# Synaptotagmin-7 endows a population of chromaffin cell vesicles with enhanced calcium sensing and delayed content release properties

**DOI:** 10.1101/704205

**Authors:** M Bendahmane, AJB Kreutzberger, A Chapman-Morales, J Philippe, N Schenk, S Zhang, V Kiessling, L Tamm, DR Giovannucci, PM Jenkins, A Anantharam

## Abstract

Synaptotagmin-7 (Syt-7) is one of two major calcium sensors for exocytosis in adrenal chromaffin cells, the other being synaptotagmin-1 (Syt-1). Despite its undoubted importance, questions remain as to the functional and physiological role of Syt-7 in secretion. We examined this issue using two distinct preparations - mouse chromaffin cells lacking endogenous Syt-7 (KO cells) and a reconstituted system employing cell-derived vesicles expressing either Syt-7 or Syt-1. First, we report using immunofluorescence that Syt-7 exhibits a punctate intracellular distribution consistent with its sorting to organelles, including dense core vesicles. We also find that the likelihood of vesicle fusion in KO cells is markedly lower than in WT cells. When fusion does occur, cargoes are discharged more rapidly when only Syt-1 is available to facilitate release. A distinctive characteristic of KO cells is that secretion runs down after prolonged cholinergic stimulation. In contrast, exocytosis persists in WT cells even with extended exposure to acetylcholine, suggesting a key role for Syt-7 in sustaining the secretory response. To determine the extent to which the aforementioned results are attributable purely to Syt-7, vesicles expressing only Syt-7 or Syt-1 were triggered to fuse on planar supported bilayers bearing plasma membrane SNARE proteins. Here, as in cells, Syt-7 confers substantially greater calcium sensitivity to vesicle fusion than Syt-1 and slows the rate at which cargos are released. Overall, this study demonstrates that by virtue of its high affinity for calcium, Syt-7 plays a central role in regulating secretory output from adrenal chromaffin cells.

## Introduction

Synaptotagmin-7 (Syt-7) belongs to a family of proteins, numbering 17 in total, many of which serve as Ca^2+^ sensors for release in a variety of secretory systems (1, 2). In the context of the adrenal medulla - a critical effector arm of the sympathetic nervous system - synaptotagmins regulate the release of adrenergic hormones and peptides from chromaffin cells (2). These released agents modulate the function of a multitude of peripheral organs, including the heart, lungs, and digestive system (3–6).

By now, it is generally acknowledged that Syt-7 and a related isoform, Syt-1, are responsible for the vast majority of Ca^2+^-dependent exocytosis from chromaffin cells (7). How-ever, there are important unresolved issues with respect to how Syt-7 operates. For example, it remains unclear as to whether Syt-7 is sorted to the plasma membrane (8) or the vesicle membrane (9, 10), and whether Syt-7 promotes (11), or alternatively, constrains (12) fusion pore expansion. It is also not obvious why two synaptotagmin isoforms are necessary to support exocytosis in chromaffin cells.

Here, we employ two distinct preparations to address questions related to the localization and function of Syt-7. The majority of the experiments in this study utilize chromaffin cells harvested from wild-type (WT) and Syt-7 knockout (KO) mice. First, we performed immunolabeling studies in dispersed mouse chromaffin cells to show that endogenous Syt-7 is primarily sorted to intracellular organelles (including dense core vesicles) rather than the plasma membrane. The role of Syt-7 in controlling the characteristics of exocytosis was inferred by monitoring - with TIRF microscopy - the discharge of fluorescently labeled lumenal cargo proteins. Using this approach, we demonstrate that cells without Syt-7 release cargos at significantly faster rates than WT cells expressing a full complement of synaptotagmins. Consistent with data from overexpression studies (12, 13), this suggests that endogenous Syt-7 constrains rather than accelerates fusion pore expansion (11).

Importantly, we find that the absence of Syt-7 severely disrupts the ability of chromaffin cells to respond to cholinergic stimulation. In fact, the likelihood of observing fusion at all declines precipitously in KO cells after the first few seconds of stimulation with acetylcholine (ACh). In contrast, WT cells expressing a full complement of synaptotagmins continue to secrete even upon prolonged exposure to ACh. The ability of Syt-7 to sustain exocytosis even in the face of rapidly declining intracellular Ca^2+^ (i.e., after nicotinic receptor desensitization) is likely due to its high Ca^2+^ affinity. This idea is underscored by reconstitution studies in which dense core vesicles consisting of only Syt-1 or Syt-7 were triggered to fuse with synthetic bilayers. Even in this reduced setting, Syt-7-bearing vesicles fuse at significantly lower Ca^2+^ concentrations than Syt-1-bearing vesicles, with fusion pores that dilate at slower rates.

Taken together, our data highlight clear functional distinctions in the properties of synaptotagmins relevant to the physiological regulation of Ca^2+^-triggered exocytosis. The data suggest that in this system, as likely in others, isoforms may have diverged to fill different niches in the physiological regulation of Ca^2+^-triggered exocytosis.

## Materials and Methods

### Animals

Litters of adult male and female Syt-7 −/− (gift of Dr. Joel Swanson; (14)) and +/+ (from a C57BL/6J background and obtained from Jackson Labs, Bar Harbor, ME) were used in these studies. Animals were group housed with 24h access to food and water, 12/12 dark/light cycle. All animal procedures and experiments were conducted in accordance with the University of Michigan Institutional Animal Care and Use Committee.

### Chromaffin cells preparation and transfection

Below, we describe a novel method for the isolation and culture of adult mouse chromaffin cells from the adrenal medulla. Although the protocol was adapted from previous studies ((15), it is different enough to warrant a more detailed description. Animals were gas anesthetized using an isoflurane drop jar and sacrificed by guillotined ecapitation (all procedures are in accordance with approved UM IACAC protocol PRO00007247). Six to eight adrenal glands/condition were rapidly isolated and moved to dishes containing ice cold dissection buffer (148 mM NaCl, 2.57 mM KCl, 2.2 mM K_2_HPO_4_•3H_2_O, 6.5 mM KH_2_PO_4_, 10 mM glucose, 5 mM HEPES free acid, 14.2 mM manitol). Under a dissection microscope, the cortex was rapidly and carefully removed using thin forceps (Dumont Swissmade, Switzerland Cat. # 72891-Bx) and thin micro scissors (World Precision Instruments, 14124-G). The isolated medullas were washed three times in three 150μL drops of enzyme solution containing (450 units/ml Papain (Worthington Biochemical #LS003126), 250 μg/ml BSA, and 75 μg/ml dithiothreitol). The medullas were then digested for 15 minutes in 0.5 ml of the enzyme solution at 37°C. After 15 minutes, the digesting solution was carefully removed and replaced by 0.5 ml of fresh enzyme solution and left for a maximum of 15 extra minutes at 37°C. The digestion was stopped by transferring the glands into an antibiotic-free culture medium (Dulbecco’s Modified Eagle’s Medium (DMEM) (ThermoFisher Scientific) supplemented with 10% Fetal Bovine Serum (FBS) (ThermoFisher Scientific)). The digested glands were then triturated by a push-pull movement through a 1 ml pipette tip (10 to 12 times). The supernatant was discarded, and the pellet re-suspended in antibiotic-free medium and triturated again in a 200 μl pipette tip for a better cell dissociation (10 - 12 times). The suspension was spun again at 1300 × g for 2.5 min. After discarding the supernatant, the pellet was re-suspended in resuspension buffer (Invitrogen, Thermofisher Scientific) for transfection. The cells were rapidly counted and desired plasmid was added (15 ng/10^6^ cells). The suspended cells were then transiently transfected by electroporation with a single pulse (1050 mV, 40 ms) using the Neon transfection system (Invitrogen, Thermofisher Scientific). In parallel, 35mm diameter dishes 14 mm glass-bottom dishes (MatTek Corporation, Ashland, MA. #P35G-1.5-14-C) were pre-coated with Matrigel (Corning, NY, Cat. #356230) diluted in DMEM (1:7) for two hours after which the dishes were washed with DMEM and let to dry. After electroporation, an antibiotic-free medium was added to cells to obtain a final concentration 1 million cells per ml. Three hundred microliters of the final solution containing the electroporated cells were then deposited in each dish. The cells were stored in an incubator (37°C, 5% CO2) for three to five hours. Culture medium with antibiotics was then added to a final volume of 2 ml (DMEM supplemented with 10% FBS, 9.52 unit/ml Penicillin, 9.52 μg/ml Streptomycin and 238 μg/ml Gentamicin (ThermoFisher Scientific)). The media was changed daily, and cells were used within 48 hours after plating. The method we describe here provides consistently healthy cells (Figure S1 which exhibit a high probability of secretion upon stimulation. Chromaffin cells were transfected with rat myc-Syt-7 or cargo proteins, including human Neuropeptide Y (NPY) and human tissue plasminogen activator (tPA). The myc-Syt-7 plasmid was a gift of Dr. Thomas Südhof. The fluorescent tag was located following the C-terminal region of the cargo proteins. NPY and tPA constructs (originally in pEGFP-N1 vectors) were provided by Dr. Ronald W. Holz.

### Western Blotting

Lysis buffer containing 8M urea, 5% SDS, and 5mM N-ethylmaleimide in water was heated to 65°C. Adrenal glands were dissected from five months-old mice and immediately frozen in liquid nitrogen. Four total adrenal glands from two mice of each genotype were dissolved into 200uL of warm lysis buffer, and homogenized using a handheld, battery-operated homogenizer. The homogenate was incubated at 65°C for 20 minutes and mixed 1:1 with 5× PAGE buffer (5% (wt/vol) SDS, 25% (wt/vol) sucrose, 50mM Tris pH 8, 5mM EDTA, and bromophenol blue). The lysates were stored at −20°C until use. Samples (10uL/well) were separated on a 4-12% NuPAGE Bis-Tris Gel in 1× NuPAGE MOPS SDS Running Buffer, for 1 hour at 175 mV. Transfer to nitrocellulose membrane occurred at 120 mV for 1.5 hours, on ice, in 1x NuPAGE transfer buffer. The membrane was blocked with blocking buffer containing 5% Bovine Serum Albumin (BSA) and 0.1% tween in TBS (TBS-T) for 1 hour, before incubation with primary antibodies (Rabbit anti Synaptotagmin-7 1:1000, Synaptic Systems, and Mouse anti alpha-tubulin, 1:10,000, Cedarlane) at 4°C, overnight. The membrane was washed 3× 15 minutes with TBS-T and incubated for 1 hour with LiCor fluorescent secondaries (1:10,000) in blocking buffer (multiplexed). After washing 3× 15 minutes in TBS-T, the membrane was imaged using LiCor Odyssey Clx imager.

### Reverse transcription and quantitative PCR

Reverse transcription was performed on mouse adrenal medullas dissected from adrenal glands and homogenized. Adrenal medullas from two animals in each WT and Syt-7 KO group are considered as one experiment. Specifically, four adrenal medullas from two animals were homogenized in one 1.5-ml Eppendorf tube on ice for ~45 seconds with motorized pestle mixer (Argos Technologies, Inc, Vernon Hills, IL). More than three experiments were performed for each target. RNeasy Mini (Qiagen, Valencia, CA) was used to isolate the RNAs. The first strand c DNA synthesis was performed with 400 ng of RNAs using the qScript cDNA SuperMix kit (Quanta Biosciences, Beverly, MA). The reverse transcription product was kept at −20°C until qPCR was performed. qPCR primers for the target genes were designed with online tools (GenScript PCR Primer Design and NCBI primer designing tool). The forward (fw) and reverse (rv) primer sequences are as follows: GAPDH fw CTGACGTGCCGCCTGGAGAA GAPDH rv CCCGGCATCGAAGGTGGAAGA Syt-1 fw GGCGCGATCTCCAGAGTGCT Syt-1 rv GCCGGCAGTAGGGACGTAGC Syt-7 fw CCAGACGCCACACGATGAGTC Syt-7 rv CCTTCCAGAAGGTCTGCATCTGG NPY fw GTGTGTTTGGGCATTCTGGC NPY rv TGTCTCAGGGCTGGATCTCT tPA fw CTCGGCCTGGGCAGACACAA tPA rv AGGCCACAGGTGGAGCATGG TH fw GCGCCGGAAGCTGATTGCAG TH rv CCGGCAGGCATGGGTAGCAT

Glyceraldehyde 3-phosphatedehydrogenase (GAPDH) was used as an endogenous control run in parallel with target genes. Each assay was performed in triplicate. For the qPCR, we used the PerfeCTa SYBR Green SuperMix (Quanta Biosciences, Beverly, MA). Ten ng of reverse transcription product was added to the master mix with 10 μM of each primer. The directions for the PCR protocol were followed per the manufacturer’s instructions. The qPCR was performed using the CFX96 TouchTM Real-Time PCR Detection System (Bio-Rad, Hercules, CA). Melting curves were analyzed to verify that no primer dimers were produced.

### TIRF microscopy for observation of exocytosis

TIRF imaging was performed using an Olympus cellTIRF-4Line microscope (Olympus, USA) equipped with a TIRF oil-immersion objective (NA 1.49) and an additional 2x lens in the emission path between the microscope and the cooled electron-multiplying charge-coupled device Camera (iXon 897; Andor Technology). The final pixel size of the images was 80 nm. Series of images were acquired at ~ 20 Hz using CellSense software with an exposure time of 30 ms and an EM gain of 100. pHl and GFP were excited using a 488 nm laser.

### Cell stimulation

All TIRF experiments were performed at room temperature ~24° C. The culture medium was replaced by pre-warmed (37°C) physiological salt solution (PSS) (145 mM NaCl, 5.6 mM KCl, 2.2 mM CaCl2, 0.5 mM MgCl2, 5.6 mM glucose, and 15 mM HEPES, pH 7.4). Chromaffin cells were individually stimulated using a needle (100-μm inner diameter) connected to a perfusion system under positive pressure ALA-VM4 (ALA Scientific Instruments, Westbury, NY). To trigger exocytosis, cells were first perfused with PSS for 5 - 10 s and then stimulated with high potassium containing solution (100 mM KCl) for 70-75 s. For the acetylcholine experiment, 100 μM acetylcholine (Sigma-Aldrich) diluted in PSS was perfused for 120 s.

### Image analysis

Fusion sites of vesicles containing GFP or pHl-tagged proteins undergoing exocytosis were identified. Regions of interest (ROIs) measuring 0.8 μm diameter were manually selected at fusion sites and image sequences were analyzed using the Time Series Analyzer v3.0 plugin on Fiji software. For each ROI, the fluorescence intensity was measured for each frame. A nearby ROI of the same size where no fusion events were observed was selected for background subtraction. Using a custom program written in Interactive Data Language (IDL; ITT, Broomfield, CO) by Dr. Daniel Axelrod (University of Michigan), background subtracted intensity versus time curves were plotted and the duration of cargo release was calculated. Briefly, a start time was determined by the user on the intensity versus time curves shortly before the rise of the fluorescence. The end time was determined as the point at which the intensity reaches its lowest level after the decline of the signal. The duration of release is obtained via fitting the curve between the start and end time with a weighted fifth degree polynomial. This method is described in details elsewhere (16). Each fusion event was further analyzed by the user to confirm that only events in which cargos were completely released were used in the analysis.

Cells transfected with GCaMP5g were analyzed to determine the relative amount of calcium influx into the cell. Three ROIs measuring 1.68 μm diameter were manually selected at different points within each cell and fluorescence was measured for each ROI for each frame. The equation ΔF/F was applied to each ROI for each frame, then the values were averaged between the three ROIs to determine the overall ΔF/F for the whole cell. The values determined were plotted versus time. The exclusion criteria used to select which cells to analyze were based on proper fixation to the dish, no response to the control stimulation, and single response tracings that indicated calcium influx in response to the stimulus trigger.

### Immunocytochemistry

Immunofluorescence imaging was performed to assess the distribution of endogenous Syt-1, Syt-7 and PAI-1 in chromaffin cells. Mouse chromaffin cells were plated on the same Matrigel-precoated 14 mm glass-bottom dishes used for TIRF imaging. All incubations and washing steps were performed on ice unless otherwise stated. Twenty-four hours after plating, the cells were fixed with 4% paraformaldehyde in phosphate buffered solution (PBS) for 30 min. The fixed cells were quickly rinsed with PBS and quenched with 50 mM NH4Cl solution in PBS for 30 min. After a brief wash from the NH4Cl solution with PBS, the cells were permeabilized with methanol for 7 min at −20° C. Following the permeabilization, the cells were washed in Tris-buffered saline (TBS) and blocked in 0.01 % gelatin solution for 30 minutes followed by another 30 min incubation in 4% donkey serum and 0.2% bovine serum albumin (BSA) prepared in TBS. Primary and secondary antibodies were diluted in TBS at 0.2%. Cells were incubated for two hours with a combination of polyclonal rabbit anti-Syt-7 antibody (Synaptic Systems, Göttingen, Germany. Cat # 105 173, 1:1200), and monoclonal mouse anti-Syt-1 (Synaptic Systems, Göttingen, Germany. Cat # 105 011, 1:1200). The cells were then washed and incubated for 70 minutes at room temperature with Alexa 488/561-conjugated anti-rabbit and anti-mouse secondary antibodies (Molecular Probes, Invitrogen). The double labeled cells were then washed and kept a 4° C until confocal imaging. For co-labeling of endogenous PAI-1 and Syt-7, both the polyclonal anti-Syt-7 and polyclonal anti-PAI-1 (Abcam cat# 66705) antibodies were made in rabbit. In this experiment, to avoid cross-labeling, the cells were first labeled with anti-Syt-7 following the protocol described above. Cells labeled for Syt-7 were then incubated for 60 minutes with polyclonal cross-adsorbed un-conjugated F(ab’)2-Goat anti-Rabbit IgG (Invitrogen, ThermoFisher, Cat. # A24539, 1:300) diluted in TBS-0.2% BSA solution to bind the possibly remaining free primary antibody sites (anti-Syt-7) not bound by the Alexa 488-conjugated Donkey anti-rabbit. Labeling for PAI-1 was performed separately. Briefly, the primary and secondary antibodies (anti-PAI1, 1:600 - Alexa 561-conjugated anti rabbit, 1:600) were incubated for 70 minutes in reaction tube pre-adsorbed in dry milk 1% diluted in TBS (17). After incubation, normal rabbit serum (Invitrogen, ThermoFisher Scientific, C at. # 016101) was added to the tube (10% volume/volume dilution) to bind the free secondary antibody for 60 minutes. The mix was then added in the Syt-7 labeled dishes for two hours. In control dishes, either the Syt-7 or PAI-1 antibodies were omitted (Figure S2) to verify the absence of cross-labeling. Cells were subsequently imaged using a Zeiss 880 confocal microscope (Zeiss, Oberkochen, Germany) with a 63× oil immersion objective in airyscan mode. Images were analyzed using Imaris Software (Bitplane, Zurich, Switzerland) using the “spot” function module for puncta detection and colocalization analysis (12). Cells were subsequently imaged using a Zeiss 880 confocal microscope (Zeiss, Oberkochen, Germany) with a 63x oil immersion objective in airyscan mode. Images were analyzed using Imaris Software (Bitplane, Zurich, Switzerland) using the “spot” function module for puncta detection and colocalization analysis (12).

### Electrophysiological recordings

Primary cultures of mouse chromaffin cells were maintained on glass bottom dishes and mounted onto the stage of a Nikon Eclipse TE2000, as previously described (12). A micro-manifold and polyamide-coated capillary was positioned in the field of view and bath solutions were exchanged through using a pressure-driven reservoir system. Standard whole-cell patch clamp methods were used record currents evoked by acetylcholine or by step depolarizations using an Axopatch 200B amplifier and Pulse Control/Igor Pro software. Patch pipettes were constructed out of 1.5 mm o.d. borosilicate glass (#TW150F-4; WPI, Sarasota, FL), coated with Sylgard elastomer and fire polished to resistances of 2.5-7 Mn. The standard intracellular recording solution contained (in mM): 128 N-methyl-d-glucamine-Cl, 40 HEPES, 10 NaCl, 4 Mg-ATP, 0.2 GTP, 0.1 Tris-EGTA, and pH adjusted to 7.2. I_Ach_ were induced by 10 s to 300 s applications of 100 μM ACh and recorded in physiological saline (125 mM NaCl, 2.5 mM KCl, 2 mM CaCl_2_, 1 mM MgCl_2_, 1.25 mM NaH_2_PO_4_, 26 mM NaHCO_3_, and 25 mM glucose, pH 7.4). For recording of I_Ca_, the superfusion solution was changed to a solution containing (in mM): 137 tetraethylammonium chloride, 5 CaCl_2_, 2 MgCl_2_, 10 HEPES, and 19 glucose, and pH adjusted to 7.2 with Tris. I_Ca_ current-voltage relationship was obtained in response to step depolarizations (30 ms) from a holding membrane potential of −90 mV. Pulses were applied in a randomized series to membrane potentials between −80 and +60 mV. All recordings were performed at room temperature.

### Materials and Methods for Single vesicle/supported membrane fusion assay

The following materials were purchased and used without further purification: porcine brain L-α-phosphatidylcholine (bPC), porcine brain L-α-phosphatidylethanolamine (bPE), porcine brain L- α-phosphatidylserine (bPS), and L- α-phosphatidylinositol (liver, bovine) (PI), and porcine brain phosphatidylinositol 4,5-bisphosphate (bPIP_2_) were from Avanti Polar Lipids; cholesterol, sodium cholate, EDTA, calcium, Opti-Prep Density Gradient Medium, sucrose, MOPS, glutamic acid potassium salt monohydrate, potassium acetate, and glycerol were from Sigma; CHAPS and DPC were from Anatrace; HEPES was from Research Products International; chloroform, ethanol, Contrad detergent, all inorganic acids and bases, and hydrogen peroxide were from Fisher Scientific. Water was purified first with deionizing and organic-free 3 filters (Virginia Water Systems) and then with a NANOpure system from Barnstead to achieve a resistivity of 18.2 MΩ/cm. Antibodies for synaptotagmin-1 (mouse monoclonal), synaptotagmin-7 (rabbit polyclonal) are from Synaptic Systems.

### PC12 Cell Culture

Pheochromocytoma cells (PC12) with endogenous synaptotagmin-1 and -9 knocked down, described previously (18), were cultured on 10-cm plastic cell culture plates at 37OC in 10% CO_2_ in 1x Dulbecco’s modified Eagle’s medium, high glucose (Gibco) supplemented with 10% horse serum (CellGro), 10% calf serum (Fe+) (HyClone), 1% penicillin/streptomycin mix, and 2 μg/ml of puromycin. Medium was changed every 2 to 3 days and cells were passed after reaching 90% confluency by incubating for 5 minutes in Hank’s balanced salt solution and replating in fresh medium. Cells were transfected by electroporation using ECM 830 Electro Square Porator (BTX). After harvesting and sedimentation, cells were suspended in a small volume of sterile cytomix electroporation buffer (120 mM KCl, 10 mM KH2PO4, 0.15 mM CaCl2, 2 mM EGTA, 20 mM HEPES-KOH, 5 mM MgCl2, 2 mM adenosine triphosphate, and 5 mM glutathione (pH 7.6), then diluted to ~14 × 10^6^/ml. Cell suspensions (~10 × 10^6^ in ~700 μl volume) and 30 μg of NPY-mRuby DNA and 30 μg of synaptotagmin-1 or -7 DNA added and placed in an electroporator cuvette with a 4-mm gap. Then two 255-V, 8-ms electroporation pulses were applied. Cells were immediately transferred to a 10-cm cell culture dish with 10 ml of normal growth medium. Cells were cultured under normal conditions for 3 days prior to fractionation.

### Dense core vesicle purification

Dense core vesicles from PC12 cells were purified as described previously (18). PC12 cells with shRNA mediated knockdowns of endogenous synaptotagmin-1 and -9 were transfected with NPY-mRuby (~20 10-cm plates) and plasmids for synaptotagmin−1 or −7 (18, 19). Cells were scraped into PBS and pelleted by centrifugation and then suspended and washed in homogenization medium (0.26 M sucrose, 5 mM MOPS, 0.2 mM EDTA) by pelleting and re-suspending. Following re-suspension in 3 ml of medium containing protease inhibitor (Roche Diagnostics), the cells were cracked open using a ball bearing homogenizer with a 0.2507-inch bore and 0.2496-inch diameter ball. The homogenate was spun at 4,000 rpm for 10 minutes at 4o C in fixed-angle micro-centrifuge to pellet nuclei and larger debris. The post-nuclear supernatant was collected and spun at 11,000 rpm (8000 × g) for 15 min at 4o C to pellet mitochondria. The post-mitochondrial supernatant was then collected, adjusted to 5 mM EDTA, and incubated 10 min on ice. A working solution of 50% Optiprep (iodixanol) (5 vol 60% Optiprep: 1 vol 0.26M sucrose, 30 mM MOPS, 1 mM EDTA) and homogenization medium were mixed to prepare solutions for discontinuous gradients in Beckman SW55 tubes: 0.5 ml of 30% iodixanol on the bottom and 3.8 ml of 14.5% iodixanol, above which 1.2 ml EDTA-adjusted supernatant was layered. Samples were spun at 45,000 rpm (190,000 × g) for 5 hrs. A clear white band at the interface between the 30% iodixanol and the 14.5% iodixanol was collected as the dense core vesicle sample. The dense core vesicle sample was then extensively dialyzed in a cassette with 10,000 kD molecular weight cutoff (24-48 h, 3 × 5L) into the fusion assay buffer (120 mM potassium glutamate, 20 mM potassium acetate, 20 mM HEPES, pH 7.4).

### Protein purification

Synataxin-1a (constructs of residues 1-288), SNAP-25, Munc18, and Munc13 (construct of residues 529-1407 containing the C1C2MUN region), and complexin-1 from Rattus norvegicus were expressed in Escherichia coli strain BL21(DE3) cells described previously (18–20).

### Formation of planar supported bilayers with reconstituted plasma membrane SNAREs

Planar supported bilayers with reconstituted plasma membrane SNAREs were prepared by Langmuir-Blodgett/vesicle fusion technique as described in previous studies (21–23). Quartz slides were cleaned by dipping in 3:1 sulfuric acid:hydrogen peroxide for 15 minutes using a Teflon holder. Slides were then rinsed in milli-Q water. The first leaflet of the bilayer was prepared by Langmuir-Blodgett transfer onto the quartz slide using a Nima 611 Langmuir-Blodgett trough (Nima, Conventry, UK) by applying the lipid mixture of 70:30:3 bPC:Chol:DPS from a chloroform solution. Solvent was allowed to evaporate for 10 minutes, the monolayer was compressed at a rate of 10 cm2/minute to reach a surface pressure of 32 mN/m. After equilibration for 5 minutes, a clean quartz slide was rapidly (200 mm/minute) dipped into the trough and slowly (5 mm/minute) withdrawn, while a computer maintained a constant surface pressure and monitored the transfer of lipids with head groups down onto the hydrophilic substrate. Plasma membrane SNARE containing proteoliposomes with a lipid composition of bPC:bPE:bPS:Chol:PI:PI(4,5)P2 (25:25:15:30:4:1) were prepared by mixing the lipids and evaporating the organic solvents under a stream of N2 gas followed by vacuum desiccation for at least 1 hour. The dried lipid films were dissolved in 25 mM sodium cholate in a buffer (20 mM HEPES, 150 mM KCl, pH 7.4) followed by the addition of an appropriate volume of synatxin-1a and SNAP-25 in their respective detergents to reach a final lipid/protein ratio of 3,000 for each protein. After 1 hour of equilibration at room temperature, the mixture was diluted below the critical micellar concentration by the addition of more buffer to the desired final volume. The sample was then dialyzed overnight against 1 L of buffer, with one buffer change after ~4 hour with Biobeads included in the dialysis buffer. To complete formation of the SNARE containing supported bilayers, proteoliposomes were incubated with the Langmuir-Blodgett monolayer with the proteoliposome lipids forming the outer leaflet of the planar supported membrane and most SNAREs oriented with their cytoplasmic domains away from the substrate and facing the bulk aqueous region. A concentration of ~77 mM total lipid in 1.2 ml total volume was used. Proteoliposomes were incubated for 1 hour and excess proteoliposomes were removed by perfusion with 5 ml of buffer (120 mM potassium glutamate, 20 mM potassium acetate (20 mM potassium sulfate was used in buffers with acridine orange labeled vesicles), 20 mM HEPES, 100 μM EDTA, pH 7.4).

### TIRF microscopy for single vesicle/supported membrane fusion assay

Dense core vesicle to planar supported bilayer fusion assay experiments were performed on a Zeiss Axio Observer 7 fluorescence microscope (Carl Zeiss), with a 63x water immersion objective (Zeiss, N.A. 0.95) and a prism-based TIRF illumination. Laser light at 514 nm from an argon ion laser (Innova 90C, Coherent), controlled through an acoustooptic modulator (Isomet), and at 640 nm from a diode laser (Cube 640, Coherent) were used as excitation sources. The characteristic penetration depths were between 90 and 130 nm. An OptoSplit (Andor Technology) was used to separate two spectral bands (540 nm - 610 nm, and 655 nm - 725 nm). Fluorescence signals were recorded by an EMCCD (iXon DV887ESC-BV, Andor Technology).

### Calcium-triggered single vesicle - planar supported membrane fusion assay

As previously described (18), planar supported bilayers containing syntaxin-1a (1-288): dodecylated (d) SNAP-25 (bulk phase-facing leaflet lipid composition of 25:25:15:30:4:1 bPC:bPE:bPS:Chol:PI:bPIP_2_) were incubated with 0.5 μM Munc18 and 2 μM complexin-1. Secretory vesicles were then injected while keeping the concentrations of Munc18 and complexin-1 constant. Dense core vesicle docking was allowed to occur for ~20 minutes before the chamber was placed on the TIRF microscope and the microscope was focused on the planar supported membrane. Fluorescent images were recorded every 200 ms while buffer containing 100 μM calcium and a soluble Alexa647 dye to monitor the arrival of calcium at the observation site was injected.

Movies were analyzed as previously described (18, 24, 25). Fusion efficiencies are reported as the percentage of vesicles in the field of view that fuse within 15 s. Fluorescent line shapes are presented as average from 20 single events as described previously (25).

## Results

### Syt-1 and Syt-7 exhibit a non-overlapping distribution in mouse chromaffin cells

Fluorescent labeling of endogenous Synaptotagmin 1 and 7 in bovine adrenal chromaffin cells revealed these proteins to exhibit a punctate intracellular distribution, with a low degree of colocalization (13). A similar pattern is observed in chromaffin cells harvested from the mouse adrenal medulla. As shown in Figure 1, endogenous Syt-1 and Syt-7 fluorescence is largely non-overlapping (in the range of 3 - 5%), punctate, and intracellular. It was previously reported that Syt-7 is sorted to the plasma membrane in PC12 cells and rat chromaffin cells (8). However, the punctate pattern of staining we observe is most consistent with the protein being sorting to organelles. The frequent co-localization with PAI-1 (Figure 1 C, D) - a dense core marker in chromaffin cells (16) - shows that Syt-7 is also sorted to secretory vesicles.

**Fig. 1.**
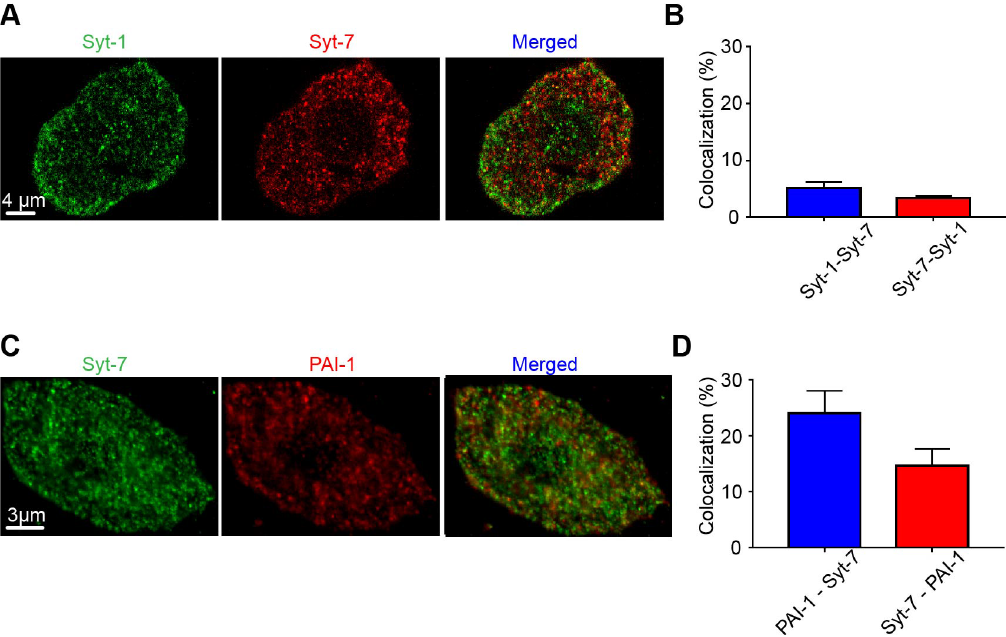
Endogenous Syt-7 is co-localized with PAI-1, a protein of the vesicle dense core. **A.** Immunolabeling was performed in WT cells for endogenous Syt-7 (red) and Syt-1 (green). Images from a representative cell showing Syt-7 fluorescence, Syt-1 fluorescence, and fluorescence of the merged channels. **B.** Bar graphs indicate the percentage of Syt-1 puncta that harbor Syt-7 (5.18 ± 1.03 %; n = 8 cells) and Syt-7 puncta that harbor Syt-1 (3.36 ± 0.43 %; n = 8 cells). **C.** Immunolabeling was performed in WT cells for Syt-7 (green) and PAI-1, a dense core vesicle marker (red). **D.** Bar graphs showing the percentage of PAI-1 puncta that are co-localized with Syt-7 (24.06 ± 1.41 %; n = 8 cells) and the percentage of Syt-7 puncta that are co-localized with PAI-1 (14.65 ± 1.075 %; n = 8 cells).

### Endogenous Syt-7 slows dense core vesicle cargo release

A major goal of this study was to delineate the ways in which the chromaffin cell secretory system depends on the presence of Syt-7. To avoid the potential ambiguities associated with overexpression of synaptotagmins in cells already expressing a full complement of endogenous proteins, most of the experiments in this study instead rely on cells harvested from Syt-7 KO mice. Figure S3A shows that compared to WT cells, expression of Syt-1 transcript is not different from WT cells, while Syt-7 transcript is almost undetectable. We also performed a Western Blot on adrenal gland lysates to verify the loss of Syt-7 protein in the KO. The alpha variant of Syt-7 (403 amino acids), targeted by Norma Andrews and colleagues when generating the original Syt-7 KO mouse, has been reported to run at approximately 45 kD (marked by arrow) (14). That band is absent in the KO (Figure S3B).

We previously showed that dense core vesicle cargo proteins NPY and Chromogranin B (CgB) were released more slowly from vesicles harboring overexpressed Syt-7 than overexpressed Syt-1 (13). Therefore, we predicted cargo release rates would be, on average, slower from WT mouse chromaffin cells than those that lacked Syt-7. To test this idea, we overexpressed NPY-pHl in WT and KO cells. Cells expressing fluorescent protein were identified by a brief exposure to 10 mM NH4Cl (26). Single cells were then depolarized by local perfusion of 100 mM KCl while exocytosis was imaged with a TIRF microscope. Figures 2 A, B, and D show that the time necessary for NPY to be completely released from fused chromaffin vesicles is broadly distributed in WT cells (times ranged from 0.049 s to 8.10 s). In Syt-7 KO cells. NPY release durations were shifted to shorter times (ranging from 0.084 s to 1.17 s) (Figure 2C and D). To confirm that Syt-7 was responsible for the slower events, rates of NPY release were also measured in KO cells co-expressing myc-tagged Syt-7 and NPY-pHluorin (myc expression was verified post hoc). In cells where Syt-7 expression was restored, NPY release times were were slowed (times ranged from 0.076 s to 8.05 s) (Figure 2D).

**Fig. 2.**
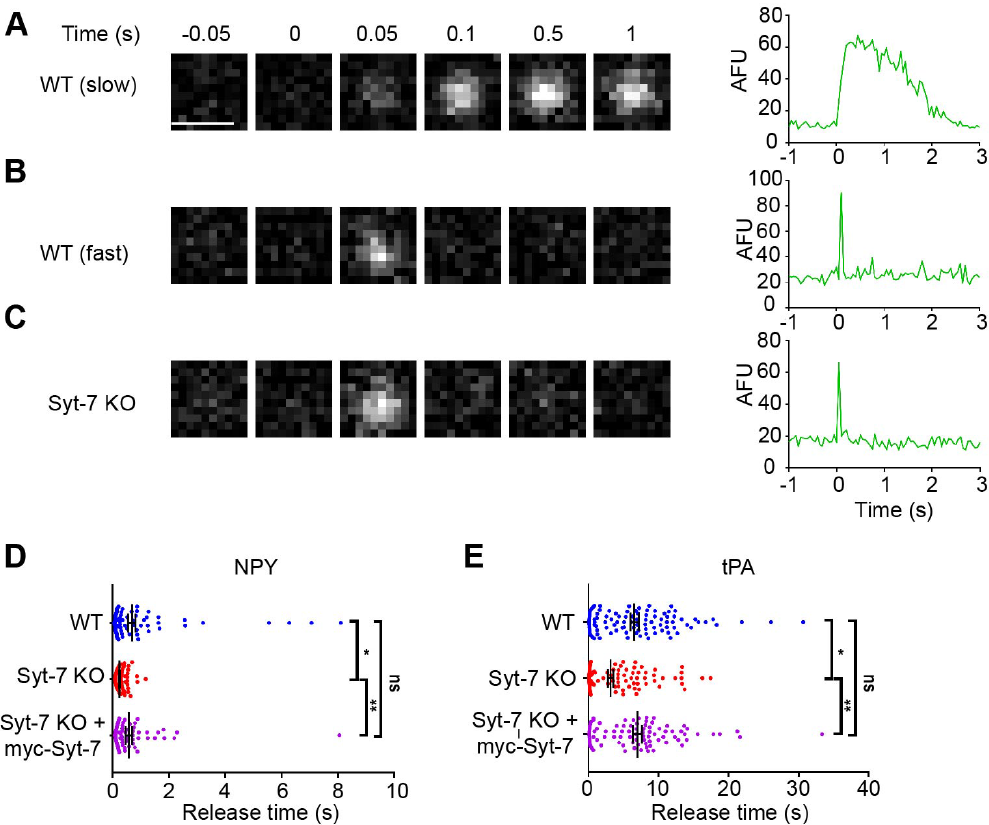
Syt-7 slows release of lumenal cargos. WT and Syt-7 KO cells overexpressing pHl-tagged NPY and tPA were stimulated with 100 mM KCl. Fusion events were observed using TIRFM and cargo release time was measured. **A-C.**. Representative images showing fusion events of NPY-pHl expressing vesicles from WT and Syt-7 KO chromaffin cells alongside intensity vs. time curves. Release times in WT cells ranged from fast to slow (A). However, release times in Syt-7 KO cells were more consistently fast (C). **D.** Release times for individual NPY events shown in a scatter plot. Release times in WT cells ranged from 49 ms to 8.10 s (average time = 0.67 ± 0.13 s; n = 100 events). In Syt-7 KO cells, the distribution of the release times was narrower, and ranged from 84 ms to 1.17 s (average time = 0.23 ± 0.01 s; n = 130 events). The overexpression of myc-Syt-7 in Syt-7 KO cells restored the broader distribution of release times, which now ranged from 76 ms to 8.05 s (average time = 0.57 ± 0.11 s; n = 77 events). **E.** Release times for individual tPA events shown in a scatter plot. Secretion events from WT cells show a large distribution of release time ranging from 68 ms to 30.67 s (average time = 6.57 ± 0.59 s; n = 103 events). In Syt-7 KO cells, the distribution of the release time was somewhat narrower, and ranged from 84 ms to 17.44 s (average time = 3.16 ± 0.38 s; n = 120 events). The overexpression of myc-Syt-7 in the Syt-7 KO cells again restored the broader distribution of release times, which now ranged from 100 ms to 33.4 s (average time = 7.00 ± 0.62 s; n = 89 events). * p<0.05, ** p<0.005 Tukey’s multiple comparison test.

Certain lumenal cargo proteins, such as tissue Plasminogen Activator (tPA), exhibit intrinsically slower release or “discharge” times during exocytosis (27, 28) In the case of tPA, this has been attributed to the protein’s ability to stabilize the curvature of the fusion pore and thereby limit its rate of expansion (16). Here, we wanted to address whether tPA release from fused vesicles may be hastened by the abrogation of Syt-7 expression - a protein which imposes its own constraints on fusion pore expansion (12, 13). As before, the rate of release of tPA-pHl was measured in both WT and Syt-7 KO cells depolarized with KCl. We found that the average rate at which tPA is released is considerably slower than it is for NPY (tPA: 6.57 ± 0.59 s, NPY: 0.67 ± 0.13 s, Mann Whitney test, p<0.0001) (Figure 2E). In fact, this is consistent with what has been reported in the bovine system by Holz and colleagues (16). As with NPY, release times for tPA were more broadly distributed in WT cells (times ranged 0.068 s to 30.67 s in WT cells) compared to Syt-7 KO cells (times ranged from 0.084 s to 17.44 s) in which, again, the slowest events were absent. Overexpression of myc-Syt-7 in Syt-7 KO cells restored these slow events. These data support the idea that Syt-7 imposes limits on the rate at which cargos are discharged during exocytosis. In the case of cargos within an intrinsically slow release profile, the effect of Syt-7 is additive.

### Vesicle fusion probability is impaired in cells lacking Syt-7

Syt-7 binds Ca^2+^ and anionic phospholipids more avidly than Syt-1. The higher in vitro Ca^2+^ sensitivity of Syt-7 (29–31) is reflected in its cellular actions. Vesicles bearing overexpressed GFP-tagged Syt-7 evince a higher fusion probability in response to KCl depolarization than vesicles bearing GFP-tagged Syt-1 (12, 13, 32). Therefore, one would predict that the fusion probability of docked vesicles in cells lacking Syt-7 would be lower than in WT cells expressing both isoforms. We tested this hypothesis by depolarizing WT and KO cells with elevated external KCl and calculating the fusion probability of docked NPY-GFP (Figure 3A). The fusion probability of vesicles in WT cells exhibited at least a 5-fold higher fusion probability than vesicles in cells lacking Syt-7 exposed to the same stimulus (Figure 3B).

**Fig. 3.**
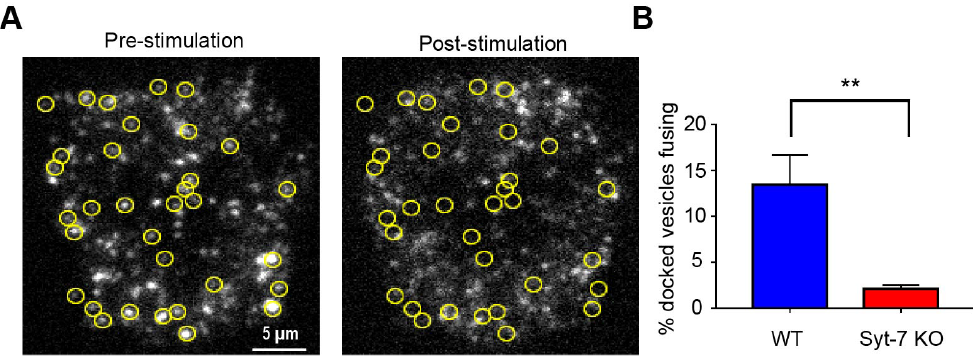
Syt-7 endows vesicles with an increased fusion probability. WT and Syt-7 KO cells overexpressing GFP-tagged NPY were stimulated with 100 mM KCl. Fusion events were observed using TIRFM. **A.** Pre- and post-stimulation images of fluorescent NPY puncta in a WT cell. Yellow circles indicate the position of vesicles that fused during the stimulation period. **B.** Bar graphs showing the percentage of docked vesicles that fused in WT (13.49 ± 3.23 %; n = 8) and Syt-7 KO cells (2.14 ± 0.36 s; n = 5). ** p<0.005, Mann-Whitney test.

We also examined another aspect of granule behavior in mouse chromaffin cells - namely, whether vesicles in KO cells exhibit the same or an altered degree of mobility than those in WT cells. These experiments were motivated by data in the bovine system showing vesicles harboring GFP-Syt-7 to be less mobile than those harboring GFP-Syt-1 (12). Here, we obtained xy tracks of vesicles only expressing endogenous Syt-1 (KO cells) and compared them to those of vesicles harboring Syt-1 and/or Syt-7 (WT cells) (Figure S4A and B). As before, NPY-GFP was used to identify vesicles. Each vesicle included in the analysis had tracks of at least 200 frames (10 s in duration). From those tracks, we calculated frame-to-frame displacement (ΔR) of individual vesicles (Figure S4C). The distribution of ΔRs for vesicles from both vesicle types were best fit to a sum of two gaussians, with the WT vesicles obviously being comprised of more than one population. The average ΔR from the slower wild type population was 29.32 +/− 1.41 nm, and 48.39 +/− 1.13 nm for the faster population, with 53.58 +/− 30.67% of vesicles residing in the slower population. The differences between the slower and faster populations of KO vesicles were much less pronounced, with a slow population ΔR of 27.82 +/− 0.59, a fast population ΔR of 39.97 +/− 3.76, and 47.84 +/− 59.77% of vesicles residing in the slower population. There is not a clear correspondence between these data and those in bovine cells in which synaptotagmin isoforms were overexpressed. However, the studies are consistent in demonstrating that chromaffin cells contain multiple populations of vesicles that evince different degrees of mobility.

### Stimulation of secretion in WT and Syt-7 KO cells with a physiological agonist

Chromaffin cell secretion, *in situ*, is triggered by activation of nicotinic receptors and subsequent Ca^2+^ influx (33, 34). Therefore, to understand how the lack of Syt-7 might disrupt secretion in a physiological setting, we stimulated WT and KO cells with ACh delivered locally via a perfusion pipet. We first measured ACh-triggered release kinetics of NPY-pHl (Figure 4A-D). As with elevated KCl stimulation, the range of NPY release times in response to ACh stimulation was broader in WT cells (0.06 s to 24.05 s) than in Syt-7 KO (0.07 s to 2.93 s) cells. On average, the rate at which NPY is released was slower in WT cells compared to Syt-7 KO cells (WT: 1.47 ± 0.34 s, KO: 0.44 ± 0.1 s, Mann Whitney test, p < 0.05) (Figure 5D). Moreover, the overall fusion probability of vesicles in Syt-7 KO cells was substantially lower than WT cells following cholinergic stimulation (3.2 ± 0.7 % KO versus 19.1 ± 3.4 %, respectively) (Figure 4A, B). One potential explanation for the difference in vesicle fusion probability observed between the two cell types is that Ca^2+^ signaling is compromised in cells lacking Syt-7. Therefore, we measured qualitative changes in intracellular Ca^2+^ in response to ACh-stimulation using the fluorescent Ca^2+^ indicator, gCaMP5g. Neither the kinetics nor the peak amplitude of the gCaMP5g signal differed between WT and Syt-7 KO cells (Figure 5C, D). We also tested the possibility that differences in the amplitude of cholinergic or Ca^2+^ currents might underlie differences in the secretory phenotype of WT and KO cells. As shown in Figures 6A-D, neither of these explanations can account for the lower fusion probability of vesicles in Syt-7 KO cells. Thus, the most parsimonious explanation is that the lack of Syt-7 protein is principally responsible for the impaired secretory response of KO chromaffin cells.

**Fig. 4.**
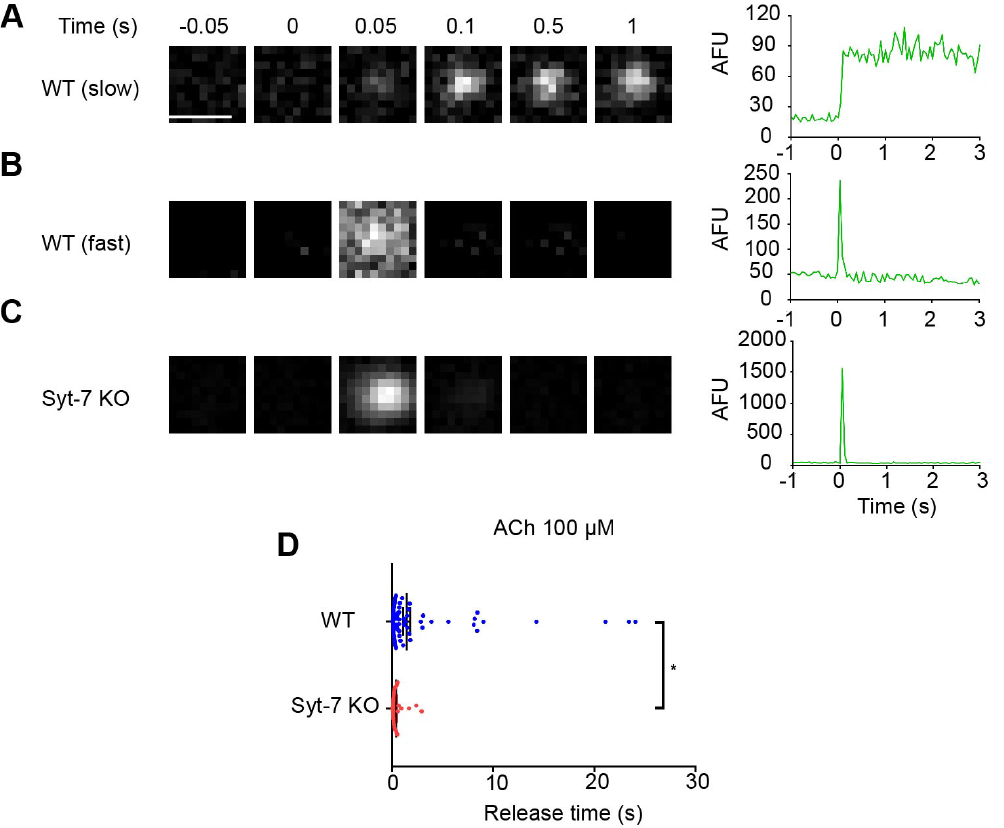
Syt-7 slows release of overexpressed NPY in cells stimulated with ACh. WT and Syt-7 KO cells overexpressing NPY were stimulated with ACh (100 μM) for two minutes. Fusion events were monitored using TIRFM and NPY release times were measured. **A-C.** Representative images showing fusion events from WT and Syt-7 KO chromaffin cells alongside intensity vs. time curves. Release times in WT cells ranged from fast to slow (A). Release times in Syt-7 KO cells were typically faster (C). **D.** Release times for individual NPY fusion events shown in a scatter plot. Release times in WT cells ranged from 67 ms to 24.05 s (average time = 1.47 ± 0.34 s; n = 130, 7 cells). In Syt-7 KO cells, the distribution of the release times was narrower and ranged from 70 ms to 2.93 s (average time = 0.44 ± 0.10 s (n = 38, 9 cells).

**Fig. 5.**
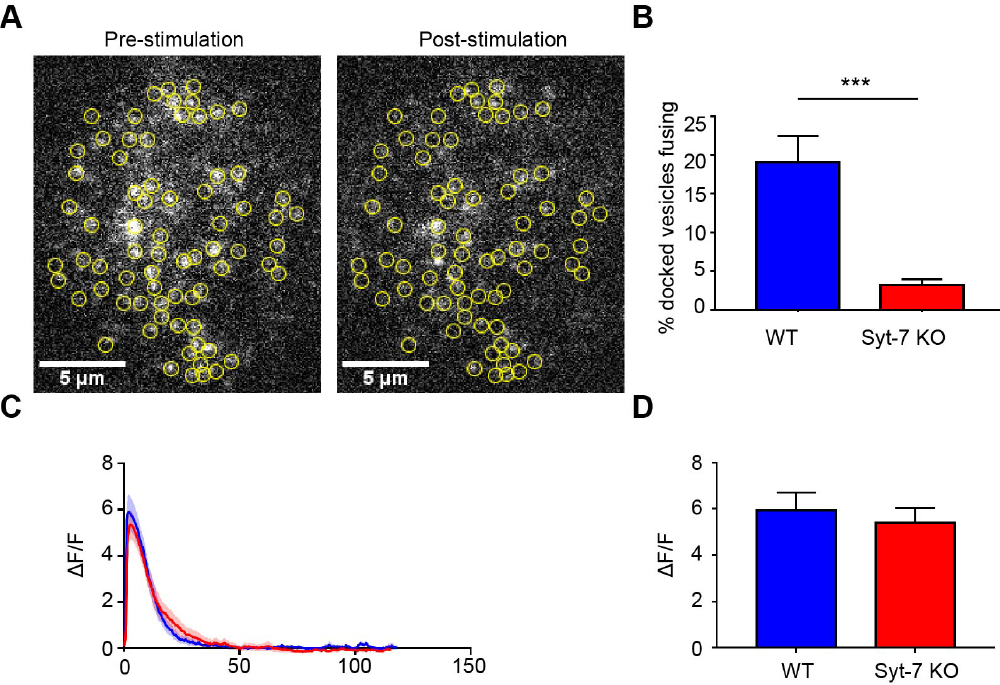
Vesicles in Syt-7 KO cells exhibit a lower fusion probability than those in WT cells in response to ACh stimulation. WT and Syt-7 KO cells overexpressing GFP-tagged NPY were stimulated with a 100 μM ACh solution. Fusion events were observed using TIRFM. **A.** Pre- and post-stimulation images of fluorescent NPY puncta in a WT cell. Yellow circles indicate the position of vesicles that fused during the stimulation period. **B.** Bar graphs showing the percentage of docked vesicles fused in WT (19.1 ± 3.4 %; n = 8) and Syt-7 KO (3.2 ± 0.7 %; n = 7). *** p<0.005, Mann-Whitney test. **C-D.** WT and Syt-7 KO cells transfected with GCaMP5g were stimulated with ACh (100 μM) for two minutes. The GCaMP signal was monitored using TIRFM, and ∆F/F was calculated for each cell. **C.** Average ∆F/F vs. time traces in response to ACh stimulation for WT (n = 7, blue) and Syt-7 KO (n= 7, red) cells. **D.** Bar graphs showing the average value of the peak ∆F/F for WT (5.41 ± 0.62) and Syt-7 KO (5.94 ± 0.76) cells. The average values were not statistically significant (Student’s t-test, p > 0.05)

**Fig. 6.**
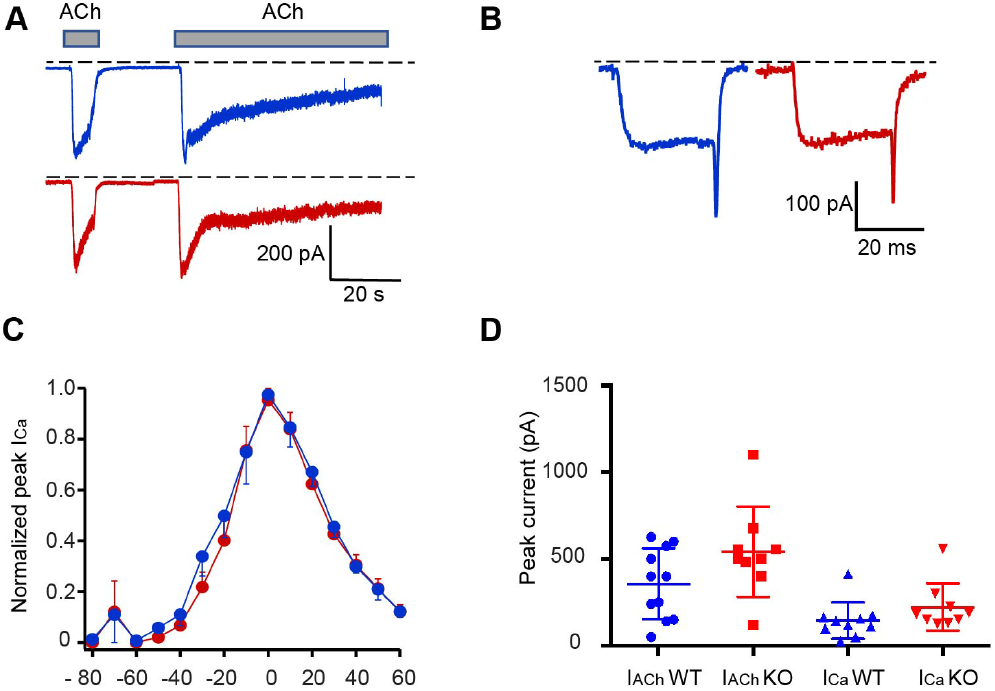
Cholinergic currents and depolarization-evoked Ca^2+^ currents are similar in WT and KO cells. Whole-cell patch clamp recordings were performed in WT (Blue) and Syt-7 KO (red) cells. **A-B.** Representative cholinergic currents (I_ACh_; A) and Ca2+ currents (I_Ca_; B). Currents were evoked by application of 100 μM ACh (A) or by a 30 ms step-depolarization (B) from a holding potential of −90 mV. **C.** Voltage-dependent activation and inactivation of the I_Ca_ in WT and Syt-7 KO cells. Each point represents the mean peak amplitude of the I_Ca_ (n = 7) evoked by a 30 ms step depolarization to test potentials of different amplitudes from a holding potential of −90 mV and normalized to the maximal response of each cell. **D.** A comparison of the mean data and range of peak amplitudes of I_Ach_ and I_Ca_ evoked in WT and Syt-7 KO cells. Mean Syt-7 KO current amplitudes were not altered compared to WT cells (Student’s t-test, p > 0.05).

### Sustained chromaffin cells ecretory output relies on the presence of Syt-7

During the course of the experiments described in Figure 4, we observed that there were kinetic differences between the fusion times of vesicles in WT and KO cells. To examine this issue more directly, we overexpressed NPY-pHl in chromaffin cells and stimulated secretion via a two minute perfusion of 100 μM ACh. The secretory response of a single WT and Syt-7 KO cell to prolonged ACh stimulation (beginning at 5 s after imaging begins) is shown in Figures 7A and B. Each of the vertical lines (blue for WT and red for KO) corresponds to the time of an individual NPY fusion event over the course of ACh exposure. In the example shown, fusion events in a WT cell continue throughout the course of stimulation. On the other hand, the secretory response rapidly wanes in the KO cell after an initial burst of fusion events. We plotted the cumulative distribution of all fusion events in WT and Syt-7 KO cells (n = 7 cells for each group) occurring after ACh exposure (Figure 7C). A distinctive feature of the cumulative time course for Syt-7 KO events is that the curve plateaus. Such a feature is consistent with the idea that Syt-7 is necessary for a sustained response to cholinergic stimulation. The cumulative time course for WT cells stimulated with ACh does not plateau; secretion persists for as long as the cell is stimulated. The curves could be fit by the sum of two exponential functions representing the fast and slow/sustained phases of secretion. The fast phase, in WT cells, is a minor component of the curve (0.66%), whereas the slow phase dominates. In the Syt-7 KO, the fast phase of release constitutes a greater proportion of the span of the curve (58.08%) than the slow phase. Interestingly, over the first approximately 20 s of ACh stimulation, there is more secretory activity in the Syt-7 KO than in the WT cell (see inset, Figure 7C). This faster phase of release may reflect the fact that Syt-1 no longer has to compete with Syt-7 for fusion sites.

**Fig. 7.**
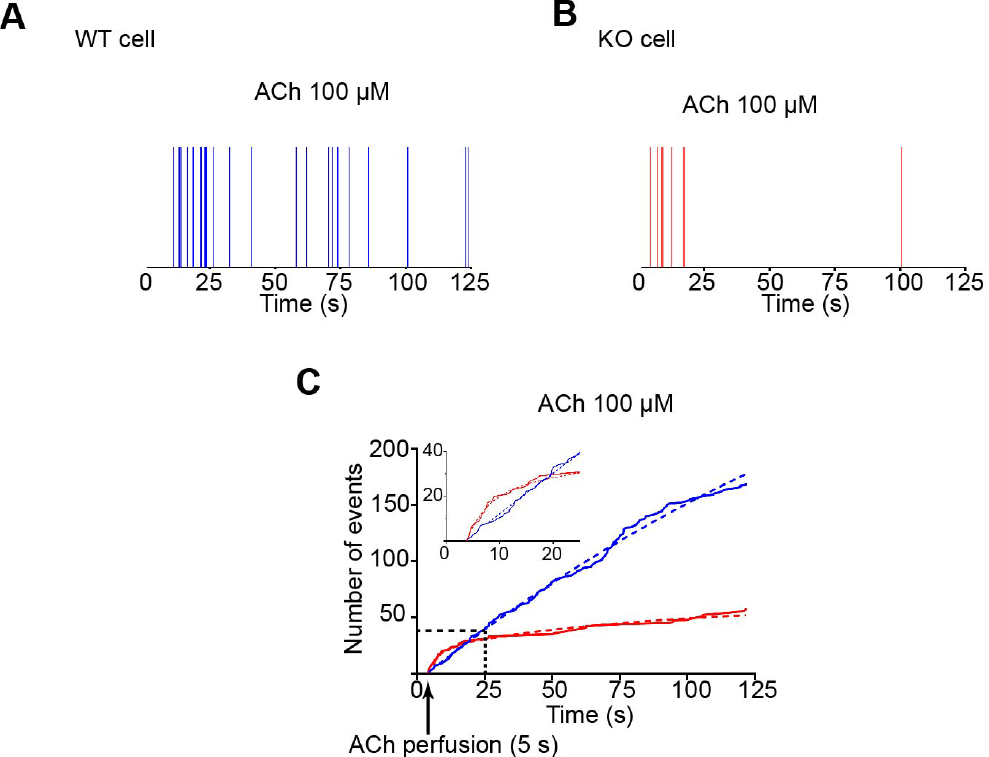
Syt-7 is necessary to sustain secretion during prolonged ACh stimulation. WNPY-pHl was overexpressed in WT and Syt-7 KO chromaffin cells. Secretion was triggered by local perfusion of 100 μM ACh. **A-B.** Time distribution of fusion events in a WT (A) and Syt-7 KO (B) cell. Each line corresponds to the occurrence of a single fusion event during the stimulation period. **C.** Cumulative frequency distribution showing the time at which fusion events occured in WT (blue) and Syt-7 KO (red) cells (n=7 for each group). WT cells continue to secrete throughout the course of the stimulation period; secretory activity in Syt-7 KO cells largely ceases after 25 sec. Note that in the 5 s - 25 s window (inset), the rate at which fusion events occur is actually higher in the Syt-7 KO cell. Curves were fit by a two phase association function in Prism 7 with R2 ≥ 0.99; WT: k_fast_ = 1.3 s^−1^; k_slow_ = 4.3 × 10−3 s^−1^; KO: k_fast_ = 0.26 s^−1^; k_slow_ = 1.3 × 10−2 s^−1^.

### Syt-7 endows dense core vesicles with distinct Ca^2+^-sensing and fusion properties

The most straightforward interpretation of our data to this point is that Syt-7 acts directly from the vesicle membrane to endow dense core vesicles with an increased fusion probability and to slow the rate of fusion pore dilation. An alternative possibility, not yet ruled out, is that the functions we have attributed to Syt-7 represent the collaborative efforts of multiple proteins on the fusion pore, with Syt-7 having a secondary role. To better assign mechanistic roles to Syt-1 and Syt-7 in exocytosis, we utilized a previously characterized purified secretory vesicle to planar supported bilayer fusion assay (Figure 8A). PC12 cells lacking endogenous synaptotagmin (18) were transfected to overexpress either Syt-1 or Syt-7 protein ((19) and Figure 8B). In the presence of SNARE regulatory proteins Munc18 and complexin-1, these vesicles will bind in an arrested state to planar supported bilayers (lipid composition of 70:30 bPC:Chol in the extracellular mimicking leaflet and 25:25:15:30:4:1 bPC:bPE:bPS:Chol:PI:PIP_2_) containing the plasma membrane SNARE proteins (syntaxin-1a and dSNAP-25). Injection of Ca^2+^ into this system readily stimulates fusion of dense core vesicles with the planar bi-layer - a process which is visualized with TIRF microscopy. Injecting increasing amounts of Ca^2+^ into the chamber containing docked dense core vesicles, triggers fusion with different efficiencies depending on the sensitivity to Ca^2+^ of the expressed synaptotagmin isoform (Figure 8C). Syt-7 containing dense core vesicles consistently fused at lower Ca^2+^ concentrations than those bearing Syt-1. The higher sensitivity of Syt-7 bearing vesicles to Ca^2+^ is consistent with the observation in cells that mild depolarization selectively activates Syt-7 containing vesicles (12, 13, 32). The intensity time course of NPY-mRuby fluorescence during release has noteworthy characteristics. Initially, a decrease in fluorescence is observed as NPY-mRuby begins to diffuse out of the early fusion pore and away from the fusion site (note dip in fluorescence in Figures 8D and 8E). After a brief delay, a sharp increase in fluorescence intensity is observed as the fusion pore expands and the vesicle membrane collapses (18). Previously, the delay time from the onset of NPY-mRuby release was shown to be sensitive to the presence of lipids with geometries that promote or inhibit fusion pore stability (25). Here, we show this feature is also sensitive to the synaptotagmin isoform expressed. A close examination of the fluorescence intensity profile of Syt-1 vesicles shows that there is approximately a 0.4 s delay time from the initial decrease in fluorescence until the collapse of the vesicle into the supported bilayer (Figure 8D). In Syt-7 vesicles, this delay is approximately 1 s (Figure 8E). The reconstituted fusion assay confirms at least two predictions based on previous studies in chromaffin cells (12, 13, 32). First, we predicted that vesicles bearing Syt-7 will fuse at lower Ca^2+^ concentrations compared to those bearing Syt-1, as the likelihood of Syt-1 or Syt-7 vesicle fusion strongly corresponded to the strength of cellular stimulation. In the in vitro reconstitution assay, we observe that Syt-7-bearing vesicles exhibit a higher fusion probability across a wide range of Ca^2+^ concentrations, up to 100 mM Ca^2+^ (Figure 8C). A second prediction, also confirmed, was that Syt-7 has the potential to impose effects on the open fusion pore that distinguish it from Syt-1. This may be because Syt-7 actively slows expansion (32), or alternatively, does not accelerate expansion to the same degree as Syt-1.

**Fig. 8.**
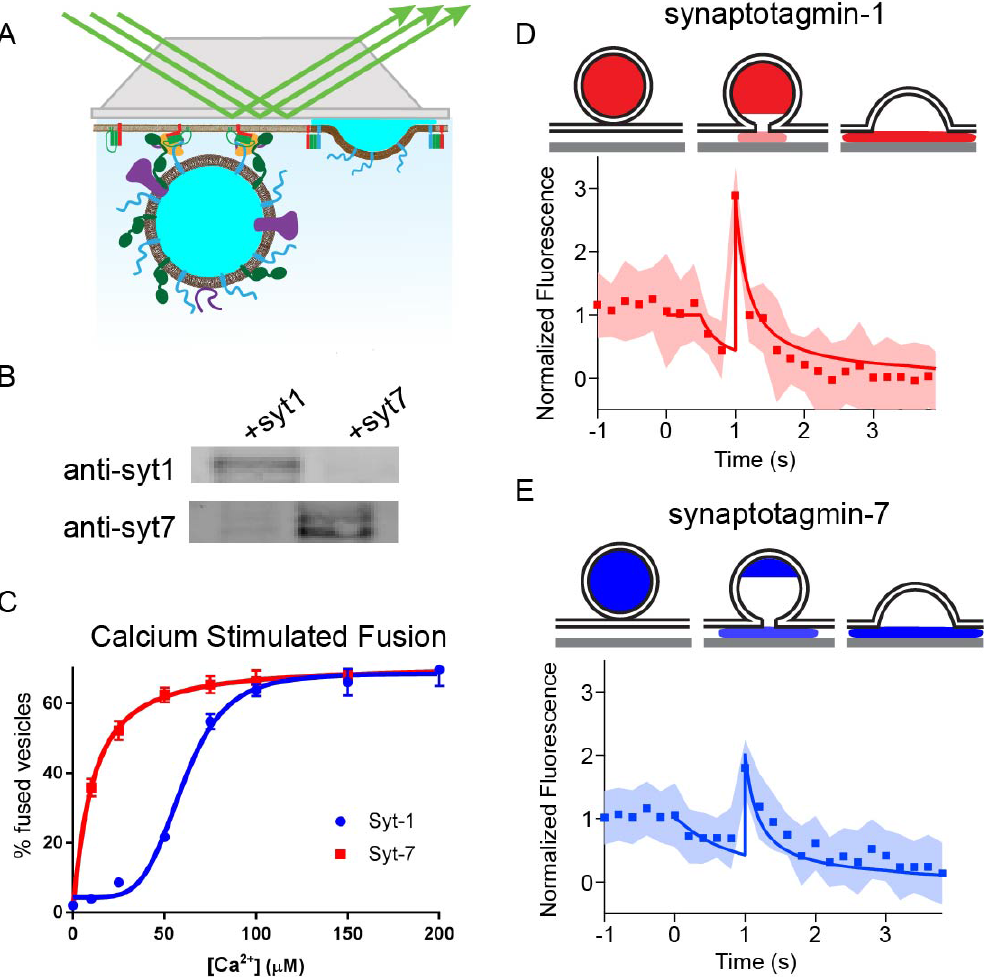
*In vitro* fusion of Syt-7 and Syt-1 vesicles on planar supported bilayers. **A-B.** Fusion of dense core vesicles containing either Syt-1 or Syt-7 with planar supported bilayers containing syntaxin-1a, dSNAP-25, Munc18, and complexin-1 were examined. As previously described, dense core vesicles labeled with NPY-mRuby stably dock to the supported bilayer, where fusion is triggered by injection of calcium. **C.** The calcium sensitivity of fusion is dependent on the Sytisoform expressed on purified dense core vesicles (K_1/2_ = 63 ± 3 μM for Syt-1 vesicles and 10 ± 1 μM for Syt-7 vesicles). Example of a characteristic NPY-mRuby signal **D-E.** is observed for fusion. There is an initial decrease in NPY-mRuby fluorescence caused by diffusion of the protein away from the site of fusion under the cleft of the supported bilayer. This is followed by an increase in fluorescence caused by a collapse of the vesicle into the supported bilayer. This action pulls the remaining lumenal content towards the brightest portion of the evanescent field. Eventually, the fluorescence diminishes as NPY-mRuby diffuses away from the site of fusion. Averaging more than 20 individual traces (D, E) for Syt-1 fusion events, revealed a short delay (D) between the initial decrease in fluorescence and the subsequent increase in fluorescence during vesicle collapse. The delay time between these phases of the fluorescent signal was prolonged for Syt-7 vesicle fusion. The cartoons illustrate the mathematical model (solid line in C and D) used to fit the data.

## Discussion

Despite the broad interest in Syt-7 as a regulator of exocytosis (1), surprisingly basic aspects of its function remain unresolved. Here, we attempted to address some of these issues in the context of the chromaffin cell secretory response. There are several important findings with respect to this goal worth emphasizing. In terms of subcellular localization, we report that endogenous Syt-7 is primarily intracellular and frequently co-localized with PAI-1, consistent with its sorting to dense core vesicles. This result is similar to what has been published by others with respect to Syt-7 localization in PC12 cells as well as primary chromaffin cells (9, 10, 35, 36). In contrast, neuronal Syt-7 has been reported to exist primarily as a plasma membrane bound protein (8, 37, 38). The reason for this discrepancy in Syt-7 localization between neurons and neuroendocrine cells is unclear. One possible scenario is that there are differences in the sorting pathways targeting Syt-7 to its eventual subcellular destination in distinct cell types. A number of possibilities exist with respect to the nature of those sorting pathways (1). Post-translational protein modifications such as glycosylation and palmitoylation have previously been shown to be important for proper sorting of some certain synaptic proteins (1, 39, 40). In fact, disrupting Syt-7 palmitoylation within macrophages abrogates its trafficking to lysosomes (41).

An unresolved question concerning the function of Syt-7 in chromaffin cells is whether it constrains, or alternatively, promotes fusion pore expansion. We previously reported that overexpressed Syt-7 stabilizes the fusion pore (13). Accordingly, vesicle cargos in Syt-7-bearing vesicles were discharged with slower kinetics than those associated with Syt-1-bearing vesicles (32). A recent study by Zhuan Zhou and colleagues posits an alternative scenario, where Syt-7 promotes rather than inhibits fusion pore expansion (11). Here, we used two distinct experimental preparations to address this issue, including primary mouse chromaffin cells lacking Syt-7 and a reconstituted single vesicle fusion assay employing vesicles only expressing one synaptotagmin isoform at a time. Both sets of experiments point to the same conclusion - that Syt-7 retards the rate at which fusion pores expand and thereby slows lumenal cargo release. We note that our findings are also consistent with TIRF-based measurements of secretion in other systems, including mouse embryonic fibroblasts, where it was shown that Syt-7 restricts post-fusion soluble content release and diffusion of vesicle membrane proteins into the plasma membrane (42). It should also be noted that multiple cellular effectors have been identified to modify fusion pore properties including some acting from the cytosol (dynamin, Epac2, myosin II etc.) (5, 43–47) and others acting from within the vesicle itself (16, 28, 48). Thus, it is very likely that Syt-7 cooperates with other proteins to modify the properties of release in ways that are not yet clear. Nevertheless, the work here points to a role for Syt-7 that is at least as important as those of the other effectors referenced above.

Much of our earlier work on the synaptotagmins has been performed in the bovine chromaffin cell system. We note here that a significant difference between the bovine and mouse preparations is that 25 mM KCl - a “mild” stimulus we previously used to evoke secretion in bovine chromaffin cells - fails to evoke secretion with any consistency in the mouse chromaffin cells (not shown). Instead, it was necessary to use 100 mM KCl to obtain an analyzable number of fusion events; this concentration of KCl was previously at the strong end of our stimulation regime (12). Note, that neither 100 mM KCl nor 100 μM ACh is as effective in evoking secretion in Syt-7 KO cells as in WT cells. The greater likelihood of observing fusion events in WT cells is not due to impaired Ca^2+^ “handling” or excitability in Syt-7 KO cells. Instead, we attribute the deficiency in the secretory response to the absence of the high affinity Ca^2+^-sensor. This hypothesis is supported by the observation that there is effectively no difference in the magnitude of Ca^2+^ or cholinergic current in WT versus KO chromaffin cells (Figure 6). Moreover, the dose-response curves in Figure 8C show that there is at least a 6-fold difference in the [Ca^2+^]_1/2_ for fusion of purified dense core vesicles bearing Syt-1 (63 μM) versus Syt-7 (10 μM). In fact, at 10 μM Ca^2+^, there is no appreciable fusion of Syt-1 vesicles.

The presence of both Syt-7 and Syt-1 is necessary for an effective secretory response to a range of intracellular Ca^2+^ levels, from low to high. This is evident from experiments showing that the ability of chromaffin cells to sustain secretory output is compromised in the absence of Syt-7 (Figure 7). In Figure 9, we propose a mechanism to explain why such an outcome might be observed. Over the course of a two-minute exposure to ACh, nicotinic receptors undergo densensitization (Figure 6A). Free cytosolic Ca^2+^, initially elevated as a result of Ca^2+^ influx through nAChRs, and also possibly Cavs, is rapidly sequestered, buffered, or extruded (49). Despite the collapse of the Ca^2+^ gradient, fusion persists in WT cells throughout the period of ACh perfusion (Figure 9B, C). On the other hand, in Syt-7 KO cells, secretory activity quickly ceases after an initial burst of fusion. Thus, we conclude that expression of Syt-7 is necessary for sustained activity in the face of rapidly declining levels of intracellular free Ca^2+^ following nAChR desensitization. Presumably, the residual Ca^2+^ that remains is effective at triggering fusion (note as well the distinct Ca^2+^ dose-response curves for fusion of dense core vesicles (Figure 8C)) are likely sufficient to account for this phenomenon.

**Fig. 9.**
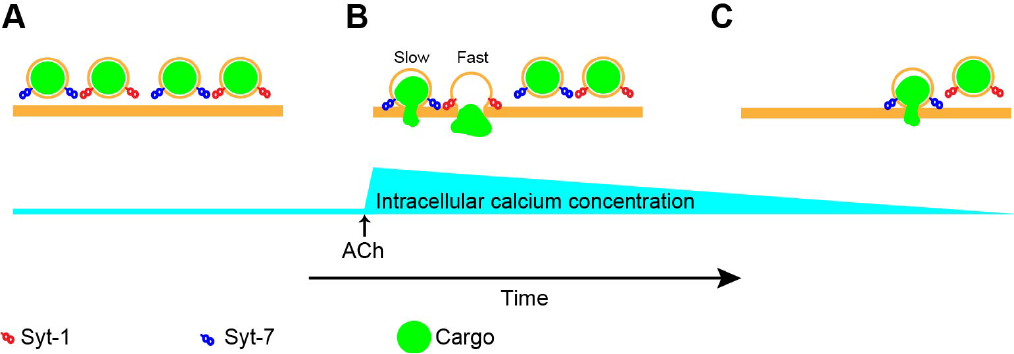
Chromaffin cells require Syt-7 for sustained secretory activity during cholinergic stimulation. **A-B.** Chromaffin cell vesicles harbor Syt-1, Syt-7, or both (not shown). When stimulated with ACh, intracellular calcium concentrations increase rapidly (Figure 5C), leading to fusion of Syt-1 and Syt-7 vesicle populations. **C.** Prolonged stimulation with ACh causes nicotinic desensitization. Associated with desensitization is a corresponding decline in intracellular Ca^2+^ which is buffered, sequestered, or otherwise removed from the cytosol (Figures 5C and 6A). Because Syt-1 has a lower affinity for Ca^2+^ than Syt-7, vesicles bearing this isoform are less likely to fuse as intracellular Ca^2+^ levels decline. On the other hand Syt-7-bearing vesicles continue to fuse even with diminished free cytosolic Ca^2+^.

The assertion that the initial and later phases of exocytosis have different Ca^2+^ sensitivities is supported by previous work by Holz and colleagues in permeabilized bovine chromaffin cells (50). By measuring [3H] norepinephrine in the medium after cells were exposed to different levels of Ca^2+^, M. A. Bittner and R. W. Holz (50) concluded that an early phase of release (i.e., occurring with 5 s of stimulation) must correspond to a lower affinity process ([Ca^2+^]_1/2_ of approximately 100 μM) than a later phase of release (i.e., occurring between 5-10 s ([Ca^2+^]_1/2_ of approximately 10 μM)). In this study, we attribute a molecular mechanism to their observation based on the sequential activation of specialized low and high affinity Ca^2+^ sensors.

To conclude, the experiments in this study demonstrate that the absence of Syt-7 has a profound impact on the Ca^2+^-triggering of secretion from chromaffin cells. It is, therefore, somewhat surprising that the global Syt-7 KO mouse does not present any overt phenotypic disturbances. At least in the context of the adrenal medulla, we would predict that there should physiological consequences to the lack of Syt-7. But it may be that, in the KO, compensatory mechanisms exist to ensure adequate delivery of hormones to peripheral organs to maintain homeostasis. Even in this scenario, it seems likely that physiological disturbances will become evident if a system without Syt-7 is challenged with the appropriate stressor. Exactly what form that stressor should take, however, is not yet established.

## ACKNOWLEDGEMENTS

The authors thank Drs. J. David Castle and Ronald W. Holz for critical reading of the manuscript. This study was supported by NIH grants R01GM111997 to A.A. and P01GM072694 to L.K.T. A.C-M. and J.M.P. are supported by a fellowship from the Pharmacological Sciences Training Program T32GM007767 at the University of Michigan.

---

**Fig. S1.**
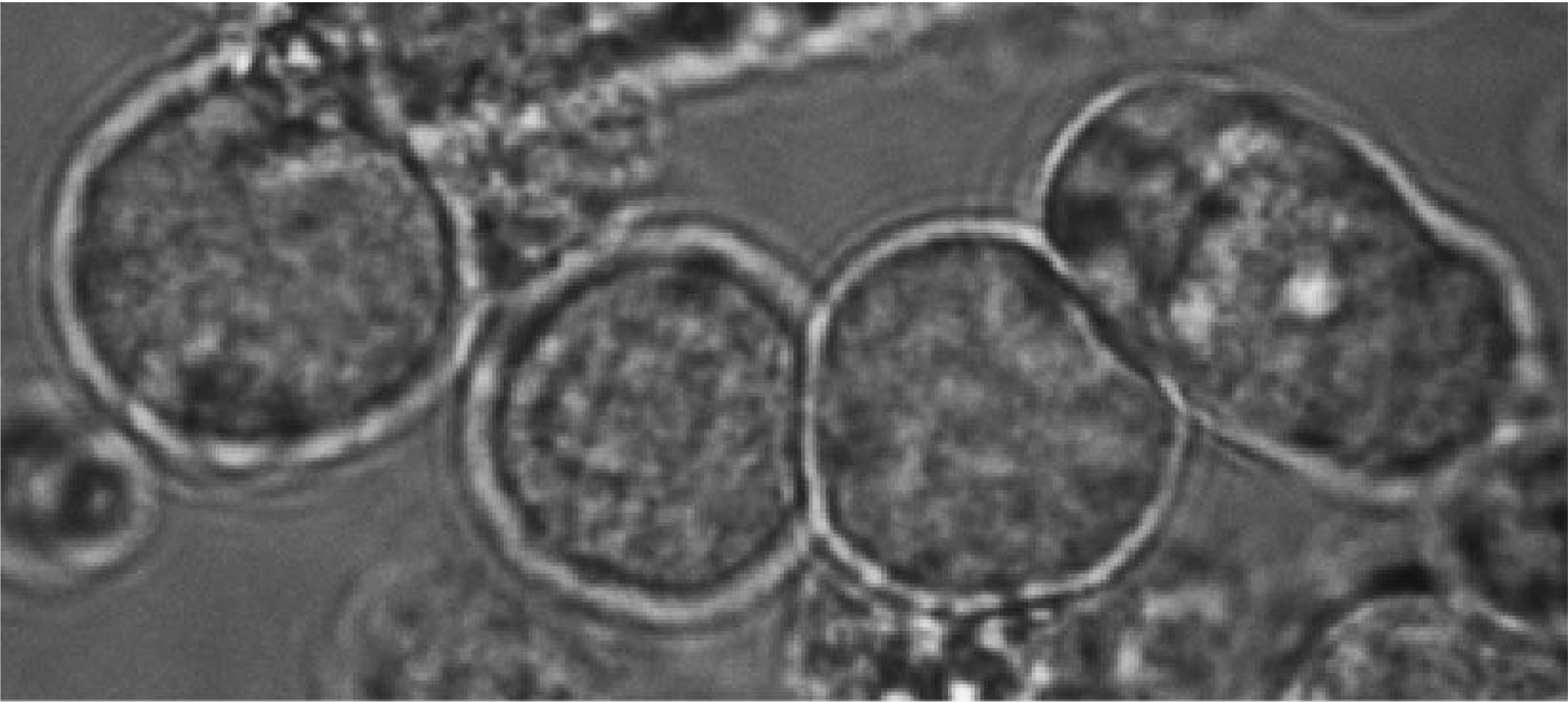
Mouse chromaffin cells 24 hours after plating on Matrigel-coated glass bottom dishes. Mouse chromaffin cells 24 hours after plating on Matrigel-coated glass bottom dishes.

**Fig. S2.**
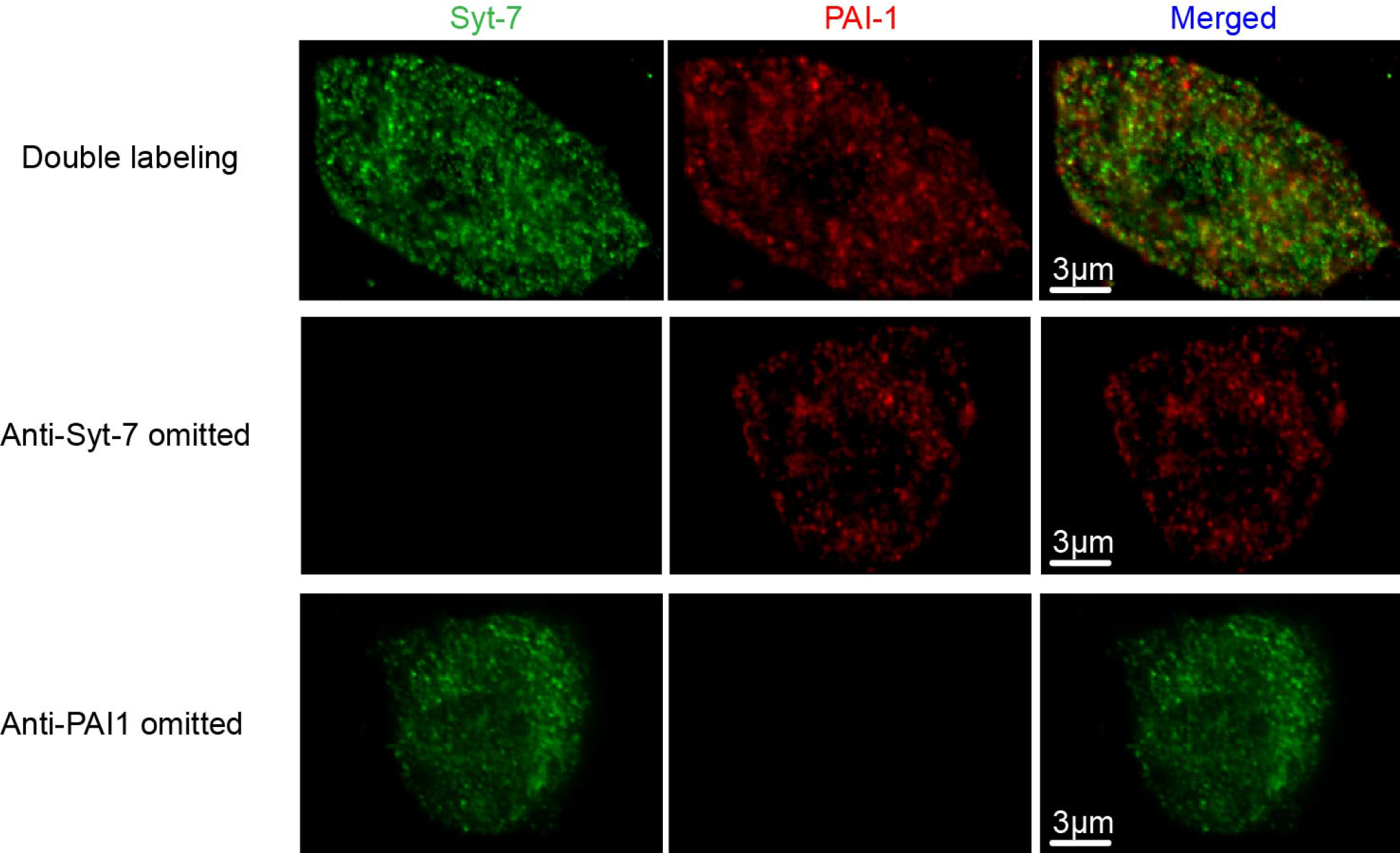
Validation of immunocytochemical-staining method using two primary antibodies raised in rabbit. Images show double-labeling for Syt-7 and PAI-1 (top row, image from figure 1). No detectable labeling was seen when of one primary antibodies was omitted (Syt-7, middle row and PAI-1 bottom row).

**Fig. S3.**
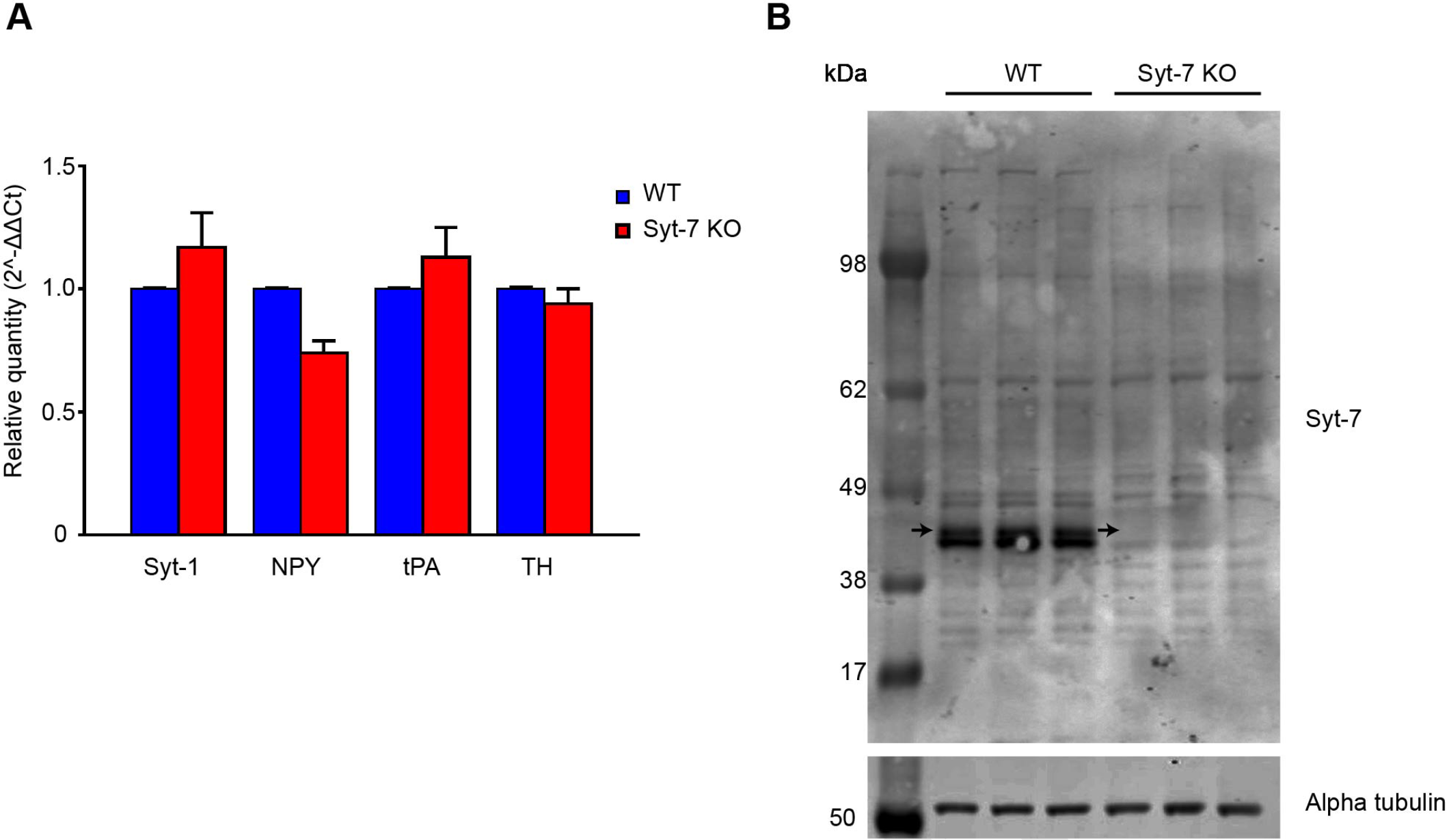
**A**. Expression of Syt-1, NPY, tPA, and TH transcripts in WT and Syt-7 KO chromaffin cells. WT and KO adrenal medullas were homogenized, mRNA was extracted and RT-qPCR was performed on the samples for Syt-1 (n = 5), NPY (n = 3), tPA (n = 3), TH (n = 3), and Syt-7 (n = 2). Syt-7 transcript was not detected using the primers shown in Table 1. Expression of Syt-1 mRNA was not significantly different in WT and KO cells (Student’s t-test, p > 0.05). **B.** Western Blot analysis was performed on WT and Syt-7 KO adrenal glands. Boxed region (white) indicates the region where the approximately 45 kDa Syt-7 alpha variant should be observed (8, 51). It is present in the WT lanes but not in the Syt-7 KO lanes.

**Fig. S4.**
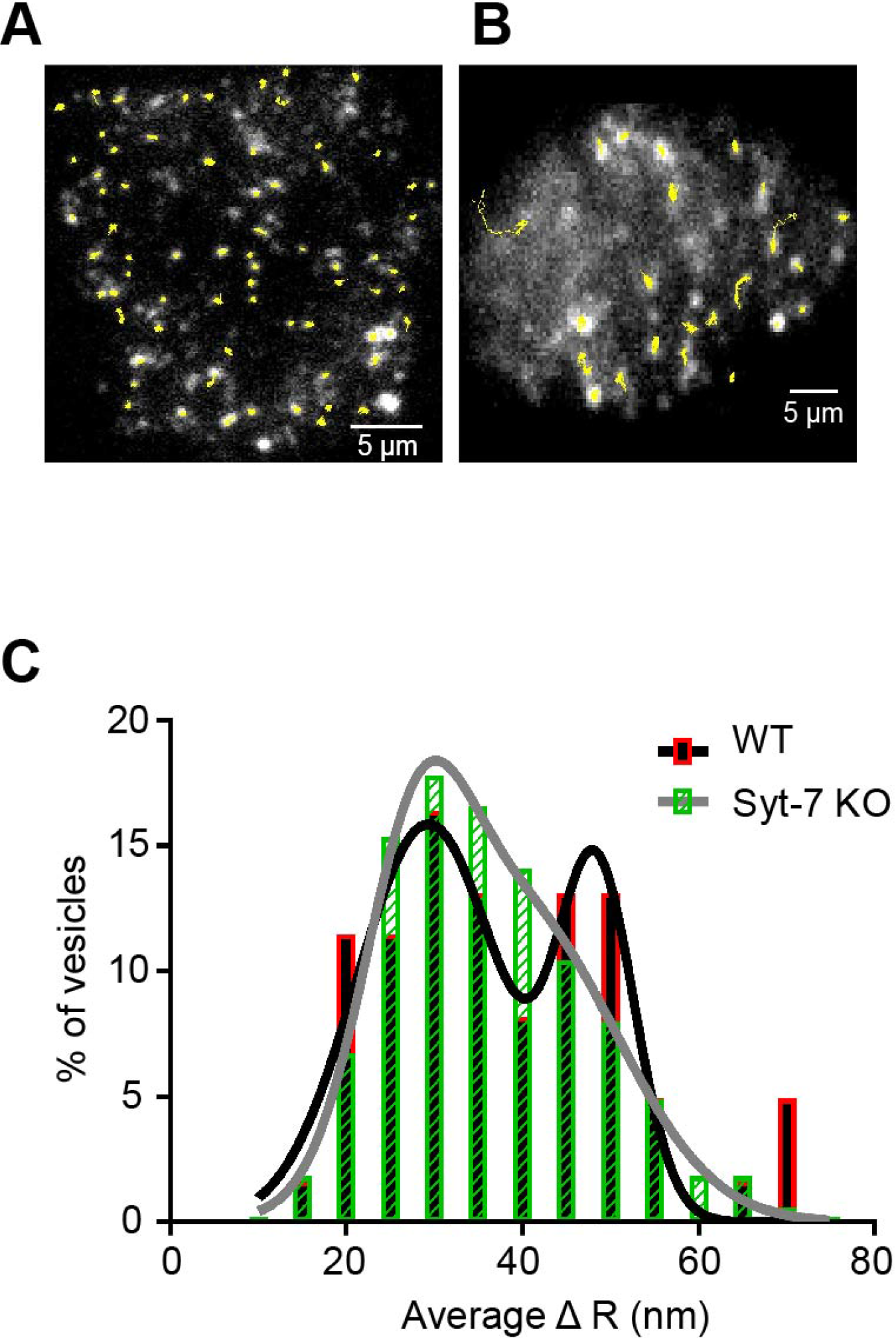
An analysis of vesicle motion in WT and Syt-7 KO cells. **A-B.** Representative tracks of WT (A) and Syt-7 KO (B) vesicles. **C.** Average mean frame-to-frame displacement of slower and faster WT vesicle populations, 29.32 +/− 1.41 nm/frame and 48.39 +/− 1.13 nm respectively, and slower and faster Syt-7 KO vesicles, 27.82 +/− 0.59 nm/frame, and 39.97 +/− 3.76 nm/frame respectfully.

**Movie S1. Example of a WT mouse chromaffin cell expressing NPY GFP and stimulated with 100 mM KCl.** Secretion was imaged with a TIRF microscope (see Methods). Corresponds to cell shown in Figure 3.

**Movie S2. Example of a WT mouse chromaffin cell expressing NPY GFP and stimulated with 100 μM ACh.** Secretion was imaged with a TIRF microscope (see Methods). Corresponds to cell shown in Figure 5.

